# Human essentiality genes and targeted oncology therapies

**DOI:** 10.1101/2020.08.18.256198

**Authors:** Andrew R. Harper, Ultan McDermott, Slavé Petrovski

## Abstract

Human essentiality genes are significantly enriched in targeted therapies successfully used in oncology. Embedding human essentiality metrics into discovery pipelines could optimise the delivery of highly effective targeted therapies among clinical development strategies.

## Main text

### Background

Cancer, a disease of the genome attributable to both inherited germline and acquired somatic variants, has proven amenable to therapeutic perturbation through the exploitation of underlying genetic vulnerabilities.^1^ As such, almost all FDA approved targeted therapies are directed against activating missense variants, inframe indels or genomic amplifications in established oncogenes. However, targeted therapies are only viable when a compatible genomic aberration is identifiable, and consequently there remains a large unmet need for highly effective, tolerable cancer drugs.^2^ The pursuit of potential therapeutic targets has benefitted from global initiatives, such as the Pan-Cancer Analysis of Whole Genomes Consortia, and large CRISPR-based dependency screens across cancer cell lines (https://depmap.org/portal/ and https://score.depmap.sanger.ac.uk/).^3^ Highlighting a panoply of putative genes involved in cancer biology, a key challenge now relates to the identification and prioritisation of suitable therapeutic targets. Minikel et al. (2020) propose genes that are intolerant of predicted loss of function (LoF) variants, otherwise known as human essentiality genes, may make good drug targets.^4^ Here, we provide perspective on how human essentiality data, derived from large cohorts of human germline exomes, can be used to help enhance current target selection pipelines specifically for oncology targeted therapies.

### Human Essentiality genes

Essentiality genes are necessary for the viability of an organism. Metrics that aim to approximate human essentiality measure a gene’s intolerance to naturally occurring LoF variants in the general population. Each metric has a specific set of underlying assumptions.^5^ These metrics include the ratio of observed-to-expected predicted loss-of-function variants (LOEUF), the probability of being LoF intolerant (pLI) and the LoF FDR (FDR p-value for preferential LoF depletion)).^6–8^ Using the LOEUF metric, genes known to harbour driver variants (median: 0.16 [IQR: 0.06-0.51]; Mann-Whitney U test *P* = 3.39 × 10^−46^), oncogenes (median: 0.12 [IQR:0.04-0.25]; Mann-Whitney U test *P* = 4.56 × 10^−24^) and tumour suppressor genes (median: 0.10 [IQR:0.04-0.23]; Mann-Whitney U test *P* = 2.45 × 10^−35^) are all significantly more LoF intolerant than the total collection of protein-coding genes (n=19,141; median: 0.48 [IQR:0.20-0.78]). Of greatest relevance to target prioritisation, genes known to be perturbed by current FDA approved targeted cancer therapies were enriched for more constrained scores than the overall protein-coding exome (median: 0.18 [IQR:0.06-0.39]; Mann-Whitney U test *P* = 4.44 × 10^−22^).

The protein-coding exome can also be dichotomised, based on the upper bound of the observed/expected ratio’s 90% confidence interval (i.e., genes with LOEUF<0.35), into high constraint versus the rest of exome. Whilst only 15.5% (n=2,968/19,141) of all human genes are germline essential based on this threshold (LOEUF<0.35), remarkably almost half (n=82/176) of the genes perturbed by successful targeted cancer therapies are among the human essentiality genes (*P*= 1.9×10^−25^; OR=4.86 [95% CI: 3.60-6.55]) (Figure 1).

**Figure 1:**
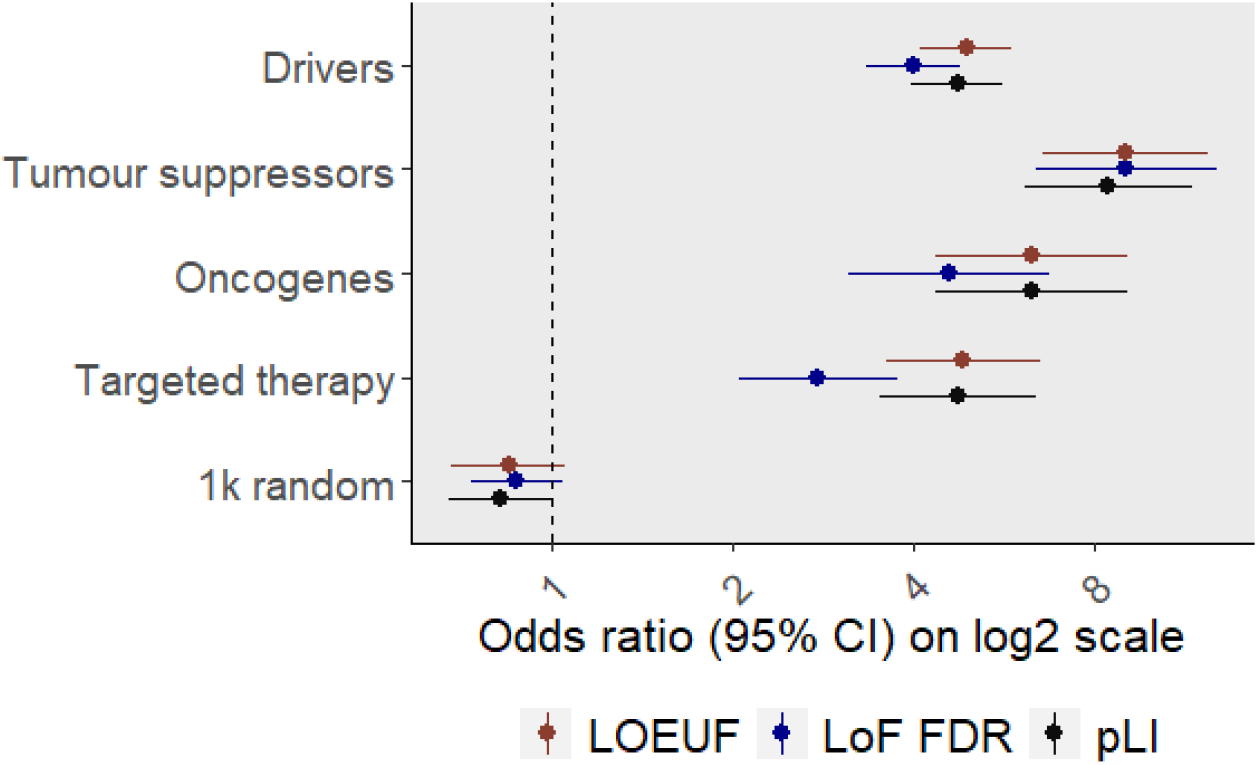
FDA approved targeted therapies for oncology appear enriched for human essentiality genes. Genes perturbed by oncology targeted therapies appear enriched for human essentiality genes, as do tumour suppressors, oncogenes and genes known to harbour driver variants. As anticipated, a random sample of 1,000 genes demonstrated no evidence of enrichment for human essentiality genes. Metrics capturing human essentiality include: the ratio of observed-to-expected predicted loss-of-function variants (LOEUF), the probability of being LoF intolerant (pLI) and the LoF FDR (FDR p-value for preferential LoF depletion). The number of genes included in each group: drivers (n=526), tumour suppressors (n=161), oncogenes (n=115) and targeted therapy (n=176). Sources: genes identified as containing a driver variant were identified from the Pan-Cancer Analysis of Whole Genomes Consortia resources (https://dcc.icgc.org/releases/PCAWG/driver_mutations); tumour suppressors and oncogenes were identified from Bailey et al. (2018)^10^; a list of targeted therapies was retrieved from the National Cancer Institute (https://www.cancer.gov), with targets corresponding to these therapies sourced from OpenTargets (https://www.opentargets.org/) and MyCancerGenome (https://www.mycancergenome.org/).

### Integration of human essentiality metrics into drug discovery pipelines

Most targeted therapies exploit the concept of oncogene addiction, whereby the disruption of a single dominant oncogene is sufficient to arrest cancer growth. As such, many oncology pipelines consult CRISPR-based cellular dependency screens during target validation. Augmenting such approaches with the germline-based human essentiality data may prove valuable. One possible concern regarding the targeting of essentiality genes relates to toxicity; given likely ubiquitous expression profiles there is scope for undesirable toxicity to normal tissue. However, this risk needs to be formally evaluated; for example, *EGFR* is an established human essentiality gene where expression is critical during embryogenesis but inhibition later in life appears well tolerated.^9^

### Summary

Genes perturbed by FDA-approved targeted oncology therapies are significantly enriched for what we now recognise are human essentiality genes. Incorporating metrics that approximate essentiality when prioritising gene targets for novel target selection may provide an opportunity to improve the selection and delivery of effective targeted therapies to patients.

## Acknowledgements

-

